# Pulsations of engineered microvessels enhance solute clearance via heterogeneous strain-dependent poroelasticity

**DOI:** 10.64898/2026.01.08.698435

**Authors:** D. Alcaide Martín, M. Fiori, S. Lorthois, A. Bancaud, J. Cacheux, Y.T. Matsunaga

## Abstract

Tissue homeostasis relies on the exchange of interstitial fluid between blood and tissues, as well as lymphatic drainage. In regions lacking conventional lymphatic vasculature, such as the central nervous system, the mechanisms governing interstitial fluid clearance remain debated, with vascular pulsatility proposed as a potential driver of interstitial fluid transport. Using a microvessel-on-chip platform that enables pulsatile pressure actuation and real-time visualization of deformation and interstitial solute dynamics in a tissue-mimetic collagen hydrogel, we investigated how pulsatile actuation regulates solute transport. Sinusoidal pressure stimulation produced a flattened but nearly symmetric deformation response in acellular lumens, whereas HUVEC-vascularized lumens exhibited pronounced asymmetry between expansion and retraction, accompanied by strain accumulation in the surrounding hydrogel. In vascularized systems, but not acellular controls, pulsatile actuation enhanced solute clearance and increased the cycle-averaged solute front velocity compared with constant-pressure actuation. To investigate the underlying mechanism, we developed a one-dimensional large-deformation poroelastic model of the vascularized bilayer. The model reproduced net directional fluid transport and showed that bilayer poromechanical heterogeneity combined with strain-dependent nonlinear permeability generates an influx–efflux asymmetry and net fluid flow. Together, these findings identify how pulsatile pressure within microvessels can generate directional interstitial fluid transport and enhance solute clearance, potentially contributing to tissue homeostasis.

## INTRODUCTION

Maintenance of tissue homeostasis requires continuous regulation of interstitial fluid (IF) volume and composition. Fluid exchange between blood and surrounding tissues is generally described by the Starling principle^1^, which relates transendothelial fluid flux to hydrostatic and oncotic pressure differences arising from the plasma–interstitial protein concentration gradient. In most tissues, this balance also involves the lymphatic system, which drains IF and returns it with lymph-borne solutes to the circulation^2,3^. Moreover, tissue deformation can generate IF movement even in the absence of a pressure gradient. Massage^4^ and exercise^5^ are associated with cyclic tissue compression and relaxation, enhancing IF drainage toward lymphatics or venules.

At smaller scales, intrinsic rhythmic deformations of the vascular system, such as those induced by the heartbeat, were also shown to promote IF circulation^6^. Some studies isolated animal arteries and characterized the amount of fluid filtration through the arterial walls under pulsations, finding out that endothelial layers modulate arterial permeability dynamically^7^ and this modulation depends on the intraluminal pressure applied^8^. Other studies investigated if arterial pulsations play a role in clearance of specific organs, such as the brain, where the routes and transport mechanisms underlying IF clearance remain controversial^9^. The blood–brain barrier, formed by tight junctions between endothelial cells, strongly restricts fluid flow and solute exchanges between blood and brain^10^. Second, the brain parenchyma lacks conventional lymphatic vessels to drain IF^11,12^. Finally, whether arteriolar pulsatility can drive bulk IF flow toward perivenous efflux routes is still debated^13–19^ largely because characterizing IF flows in the brain remains technically challenging^20^. These limitations motivate the use of controlled *in vitro* systems to investigate how mechanical stimulation generated within the vascular system affects solute transport in biological tissues.

Enhancement of solute transport by pulsating loads has been studied in simpler soft and permeable tissues. For example, cartilage naturally experiences physiological mechanical stimulation during walking and enhanced dispersion of large solutes has been consistently reported ^21–26^. Analytical^27^ and numerical^28^ models attribute this enhanced dispersion to mechanisms that are fundamentally distinct from molecular diffusion. First, analogous to Taylor dispersion in a tube^29,30^, the velocity profile in the pores of the tissue spreads concentration fronts along the flow direction, far faster than diffusion. Second, the pore morphology introduces variability in fluid streamlines, bringing into proximity fluid with different concentrations, and inducing longitudinal and transverse dispersion^31^. However, translating these findings to pulsation-enhanced IF transport is challenging because vascular interfaces are composite structures in which an endothelial barrier is mechanically coupled to a soft tissue matrix. Under periodic pressure actuation, transport cannot be assumed to arise from a linear superposition of barrier permeability and tissue elasticity, because nonlinear coupling between these components may dominate.

We therefore developed a platform that allows simultaneous measurement of vascular deformation and interstitial solute transport under controlled pulsatile or constant pressure actuation. This platform can be used either as an acellular scaffold or with Human Umbilical Vein Endothelial Cells (HUVEC)-vascularized microvessels (MVs). We show that endothelial barriers transform symmetric pressure oscillations into asymmetric tissue deformation and cycle-averaged directional solute transport, an effect not observed in acellular matrices. To mechanistically interpret these observations, we develop 1D large-deformation poroelastic simulations of soft, permeable tissues and barriers, which suggest that the presence of the endothelial barrier combined with strain-dependent permeability is required to produce a non-zero bulk fluid flow that in turn drives directional solute clearance. Together, this study identifies a vascular-driven mechanism by which endothelial barrier poromechanics convert symmetric pressure pulsations into asymmetric interstitial flow, thereby promoting directional solute transport.

## RESULTS

### Pulsatile IF platform design and operation

The Pulsatile IF platform (PIF) design was inspired by standard 3D MV systems fabricated with 200-µm-diameter acupuncture needles^32,33^. We manufactured three parallel, co-planar lumens (500 µm apart) by laser ablation of two silicone sheets (see Methods), then bonded them by hand after oxygen-plasma surface activation (Fig. 1A-B and Sup. Fig. 1A–D). Next, we punched the assembly of silicone sheets to create the main cylindrical chamber (5 mm diameter) and six lateral ports of 3 mm (Fig. 1B). With needles positioned in the device, we poured liquid collagen into the main chamber. After gelation, we removed the needles and filled the PIF platform with buffer. The platform was used either in acellular settings or seeding the two lateral lumens with HUVECs to produce MVs (Fig. 1C and Sup. Fig. 1E). The central channel remained acellular (Fig. 1A, lower panels) to enable controlled injection of fluorescent tracers into the extracellular matrix.

**Figure 1.**
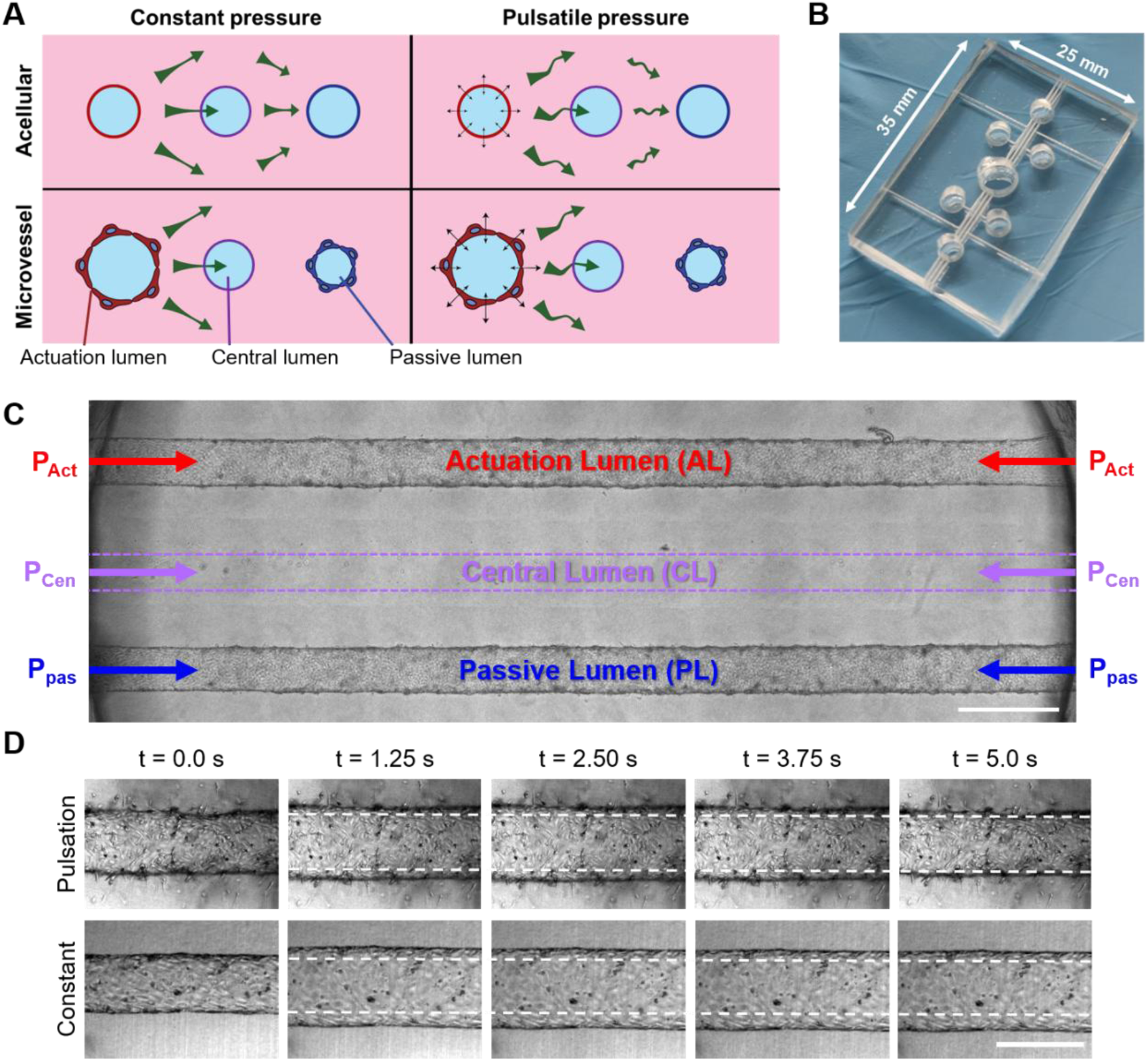
Pulsatile Interstitial Flow-on-chip (PIF chip). **(A)** Cross-section of the acellular or vascularized system with constant or pulsatile pressure actuation, the green arrows indicate the interstitial fluid flow. **(B)** Photograph of the PIF chip with its dimensions. **(C)** Bright-field micrograph of a vascularized PIF chip. MVs are fabricated in the lateral channels, while the central channel remains acellular for tracer injection. Scale bar = 400 µm. **(D)** Bright-field snippets of an MV under pulsatile pressure (top panels) and constant pressure (bottom panels). Dashed lines indicate MV diameter at t = 0 s. Scale bar = 300 µm.

Pressure regulation within each lumen was achieved using a custom 3D-printed device placed on top of the silicone chip (Methods and Suppl. Fig. 2), which allowed us to independently control the pressure in the three channels as well as on top of the collagen chamber. We imposed a higher pressure in a lateral channel, referred to as the actuation lumen (AL), while the opposite lateral channel is referred to as the passive lumen (PL), and the central lumen as CL (Fig. 1C). The pressure settings were typically defined by a baseline pressure of 100 Pa in PL, CL and on top of the chamber, and either a sinusoidal pressure input (5 s period, 100–900 Pa) or a constant pressure of 500 Pa in AL. The selected pulse period is larger than the typical heartbeat but within reported sleep-related low-frequency (0.1-1 Hz) arterial pulsation dynamics in animal models^16,17^, and slow pulsations ensure faithful actuation by the digital pressure actuator. Inspecting AL deformation by bright field microscopy showed that pulsatile pressure actuation induced periodic diameter changes of AL, whereas constant pressure expectedly maintained the actuating channels expanded (Fig 1D).

### Microvessels localize interstitial pore pressure gradients and strain

To investigate whether endothelial cells influence static lumen deformation, we measured the pressure-induced deformation of a single acellular lumen, used as the reference response of the scaffold. We recorded confocal stacks and extracted lumen cross-sections at intraluminal pressures ranging from 100 to 300 Pa, with the upper gel surface maintained at 100 Pa (Fig. 2A; see Methods). A pressure difference of 100 Pa produced a vertical strain of ∼10%, indicating an elastic modulus on the order of 1 kPa. The lumen contour was anisotropic, expanding more in the vertical than in the horizontal direction, as shown in Fig. 2A and by the horizontal and vertical axes obtained from elliptical fits to the lumen contour (Fig. 2B). This anisotropic deformation can be explained by the poromechanical response of the gel (Fig. 2C). In this single-channel configuration, fluid injection generates an interstitial pore pressure gradient oriented from the lumen to the top surface of the collagen gel, driving bulk gel deformation predominantly in the vertical direction. Furthermore, the gel was more deformable at lower pressures (red curve, Fig. 2B), consistent with the strain-stiffening behavior previously reported for collagen gels^34,35^. We fitted this static deformation using finite-element simulations based on a poroelastic model in COMSOL (see Methods), with Gent hyperelasticity as the constitutive law for the solid phase, which assumes an exponential increase in elasticity with strain (black lines in Fig. 2B top), yielding a shear modulus of 250 Pa, a limiting extensibility parameter of 0.0085, and a compressibility of 1.5 kPa. Notably, reproducing the anisotropy between the horizontal and vertical axes required accounting for the compressibility of the gel, which allows preferential swelling in the direction of the pore pressure gradient. The corresponding Poisson’s ratio was 0.33 ± 0.03 (Fig. 2D).

**Figure 2.**
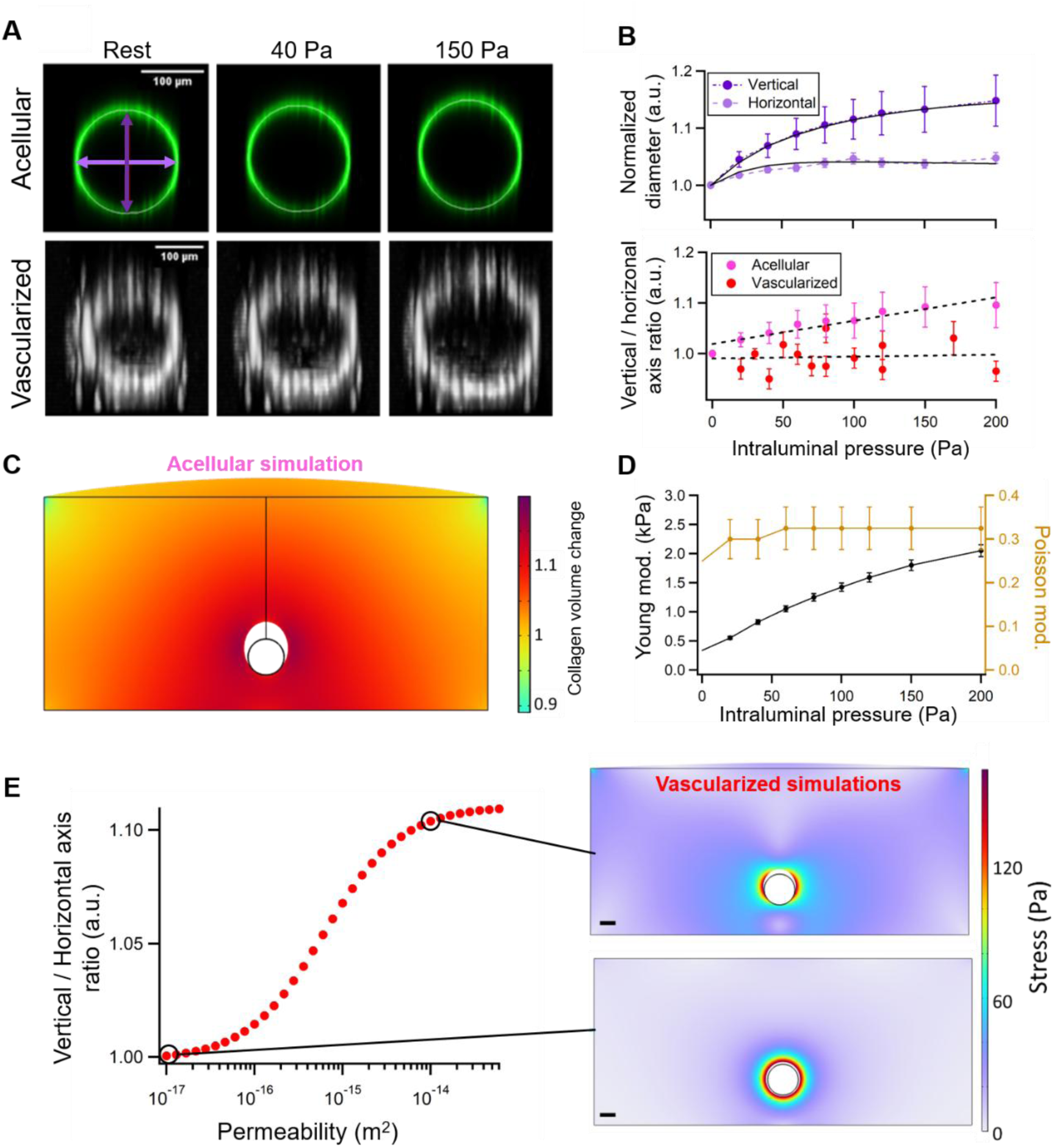
Single lumen deformation in static conditions. **(A)** Cross-section of an acellular collagen scaffold and a MV for different intraluminal pressures (upper and lower panels, respectively). The collagen gel interface is labeled with 200 nm nanoparticles and the cell monolayer with the DNA-intercalating dye Hoechst. **(B)** Top - Variation in the vertical and horizontal axes of the acellular channel as a function of intraluminal pressure (dark and light violet datasets, respectively; see arrows in A). Black solid lines: fits from numerical simulations using the Gent porohyperelastic model. Bottom - Vertical to horizontal axis ratio *vs.* intraluminal pressure for acellular scaffolds and for MVs (pink and red datasets, respectively) Dashed lines: linear fits **(C)** Finite element simulations of poroelastic deformation showing the solid volume change for an acellular lumen. Intraluminal pressure is set to 200 Pa. **(D)** Inferred Young’s modulus and Poisson’s ratio vs luminal pressure from simulations akin to that shown in Panel C. Poisson modulus remained constant around 0.3. **(E)** Parametric finite element simulations of the vertical-to-horizontal axis ratio as a function of the vascular layer permeability, using linear poroelasticity finite element simulations of vascularized lumen. Representative snapshots for permeability of 5 10^-15^ (top) and 10^-17^ m^2^ (bottom) are shown on the right. Scale bars = 100 µm.

We next experimentally investigated the expansion of a single MV and found that it maintained an approximately circular shape as intraluminal pressure increased, as shown by confocal micrographs (Fig. 2A) and by plotting the vertical-to-horizontal axis ratio as a function of pressure (Fig. 2B bottom). We attributed this shape transition to the barrier function of the endothelial cell monolayer, which localizes the pressure gradient within the endothelial layer and thereby reduces the pore pressure transmitted to the collagen gel. To test this interpretation, we performed parametric poroelastic simulations in the linear regime by varying the endothelial layer permeability. We excluded the hyperelastic response because it led to numerical instability, and used a Young’s modulus *E_gel_* = 1 kPa and a permeability *κ_gel_* = 3 · 10^−1^^4^ m^2^ for the gel, as determined previously^35^. The endothelial barrier Young’s modulus was set to *E_MV_* = 1 kPa based on indentation-based measurements, which showed that the shear modulus of endothelial cells spanned 0.5-40 kPa^36–39^. These simulations reproduced the circularization of the lumen, showing a decrease in the degree of anisotropy with decreasing endothelial layer permeability (Fig. 2E). A circular shape was reached for a permeability of *κ_MV_*∼ 10^−1^^7^ m^2^, a value that appeared reasonable for HUVEC-based endothelial barriers^40^. Furthermore, these simulations revealed stress localization within the endothelial barrier for sufficiently low permeability (snapshot in Fig. 2E). The pore pressure gradient is thereby confined within the endothelial barrier, implying that solid matrix deformation is governed by conventional, short-range elastic deformation. The endothelial barrier therefore confines the interstitial pore-pressure gradient and concentrates the resulting mechanical stress within the barrier, thereby limiting deformation of the surrounding collagen matrix.

### MVs exhibit asymmetric expansion-retraction cycles under pulsatile actuation

We next examined how pulsatile actuation affects lumen deformation in both acellular and vascularized PIF chips (Suppl. Video 1,2). Visually, AL and CL are the most affected by the sinusoidal actuation, with PL not experiencing much changes, so our analysis focused on the AL-CL region. We tracked the diameters of both AL and CL as well as the collagen thickness between them to gain localized information about the effects of pulsatile pressure on the system (Fig. 3A). Starting with AL deformation, on average, vascularized ALs exhibited larger expanded and retracted diameters than acellular ALs (Fig. 3B), with a greater deformation amplitude (35 ± 14 vs. 23 ± 7 µm for vascularized and acellular PIFs, respectively).

**Figure 3.**
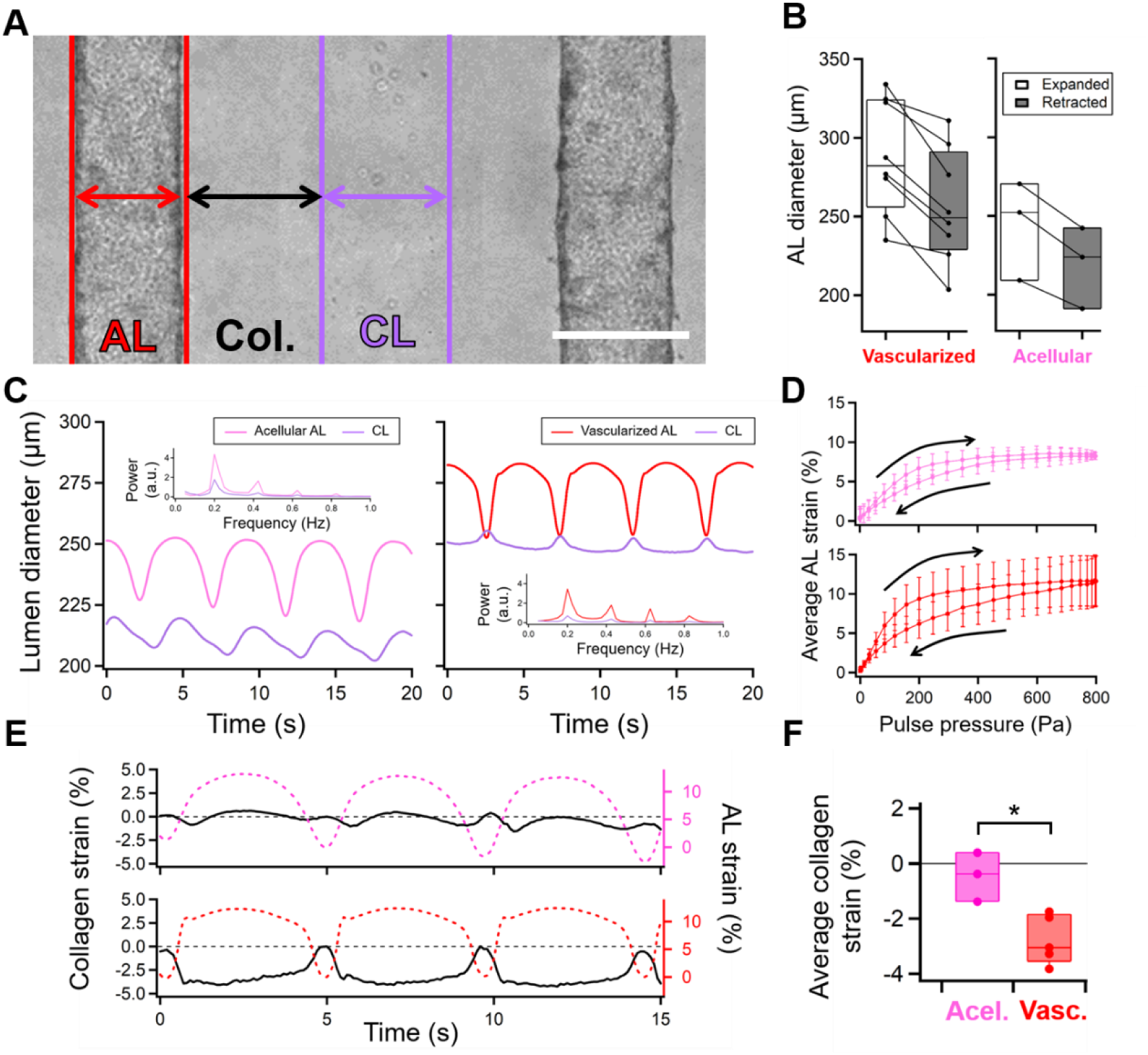
Deformation of acellular and vascularized lumens in PIF. **(A)** Depiction of the diameter-tracking strategy for the AL (red lines) and CL (purple line) in a vascularized PIF device under bright-field microscopy. Scale bar = 300 µm. **(B)** Average expanded and retracted diameters of the AL in acellular and vascularized PIFs. **(C)** Acellular and vascularized AL-CL diameter response under pulsatile pressure. Fourier transform of each signal is embedded within each graph. The intensity of high-order harmonics is greater in vascularized AL response. **(D)** Average AL strain response to a pressure cycle in acellular and vascularized PIFs. Arrows indicate strain during expansion and retraction. **(E)** Collagen gel strain obtained from inter-lumen distance variation (black curves) under pulsatile actuation: acellular PIF (top) and vascularized PIF (bottom). Dashed curves represent AL strain over time, as deduced from the data in panel C. **(F)** Cycle-averaged collagen strain in acellular (n = 3) and vascularized (n=5) PIFs (p < 0.05).

In acellular conditions, the AL exhibited deformation cycles that deviated markedly from the sinusoidal input waveform. As shown by the pink curve in Fig. 3C, ALs displayed rapid expansion and retraction phases interspersed with periods of minimal diameter variation, producing a waveform akin to a flattened sine wave. This behavior is consistent with the hyperelastic properties of the collagen gel: the material remains compliant under low intraluminal pressure but stiffens as the load increases. Fourier analysis corroborated this observation, revealing high-order harmonics in the deformation signal (subgraph in Fig. 3C left). The AL nonlinear deformation response was even more pronounced in vascularized PIFs, as evidenced by plateaus interrupted by brief, rapid diameter changes (red curve, Fig. 3C). Fourier analysis of this response revealed a power spectrum dominated by high-order harmonics, with substantially larger amplitudes than those observed in acellular lumens (subgraph in Fig. 3C right).

Further analysis of the periodic AL deformation dynamics in all acellular (n = 3) and vascularized (n = 5) PIFs highlighted an additional difference: acellular lumens exhibited nearly symmetrical expansion-retraction cycles, with only a 25% difference between the slopes of expansion and contraction (top graph, Fig. 3D), likely due to asymmetric tension/compression properties of the collagen gel ^35^. In contrast, MVs displayed marked asymmetry, with expansion occurring 61% faster than contraction (bottom graph, Fig. 3D). Fixation of vascularized samples (see Methods) increased the apparent stiffness of the pressure–deformation response (Suppl. Fig 3), and greatly decreased the non-linearities, as shown by the dominant peak at 0.2 Hz in the Fourier spectrum, consistent with the quasi-sinusoidal waveform. Altogether these findings identify expansion– contraction asymmetry and pronounced porohyperelasticity as hallmark features of live MVs.

### MV pulsations induce strain accumulation and transient release in the gel

We next investigated the propagation of the deformation within the chip by measuring the response of the CL and PL. We first observed that the CL diameter relative to the AL diameter exhibited distinct phase relationships, with a phase-shift in acellular scaffolds and phase opposition in vascularized PIFs (purple curves in the insets of Fig. 3C). The near-synchronous and opposite response observed in vascularized PIF was consistent with an instantaneous elastic-like deformation, whereas the shifted response under acellular conditions was associated with the slow propagation of interstitial pore pressure ^41^. Further, we tracked the gel deformation between the AL-CL (black arrows in Fig. 3A). Acellular PIFs displayed a global translation of the gel slab (Fig. 3E, top panel), with minimal responses on the normalized distance variations, from now on the collagen strain. Cycle-averaged collagen gel strain only amounted to −0.3 ± 0.7% (Fig. 3F). Conversely, vascularized PIFs exhibited strain accumulation during most of each pulsation cycle, only relaxing during AL retraction (Fig. 3E, bottom panel). Collagen gel strain cycle average supported this observation, with an average offset of the strain of −2.0 ± 1.3% (Fig. 3F). Notably, performing the same analysis between the AL and the PL showed greater AL-CL strain accumulation only in vascularized PIFs (Suppl. Fig. 4), likely because of MV-induced stress localization. Thus, sinusoidal pressure stimulation primarily induces translational displacement without substantial net deformation of the gel in acellular PIFs, whereas it triggers strain accumulation in vascularized PIFs, interrupted by brief relaxation phases near the minimum of the pressure input waveform.

### MV pulsations enhance solute transport

To investigate how intraluminal pulsations affect solute transport, we injected a high molecular-weight tracer (70 kDa; diffusion coefficient in collagen gels of 52 ± 7 µm²/s ^42^, Supp. Fig. 5A-C) into the CL of acellular and vascularized PIFs, and simultaneously recorded fluorescent and bright-field signals to track solute concentration and lumen deformation for 1-2 min. Kymographs (Fig. 4A) revealed broader spatial displacements in acellular settings than in vascularized PIFs, consistent with greater interstitial fluid flow in the absence of the endothelial barrier.

**Figure 4.**
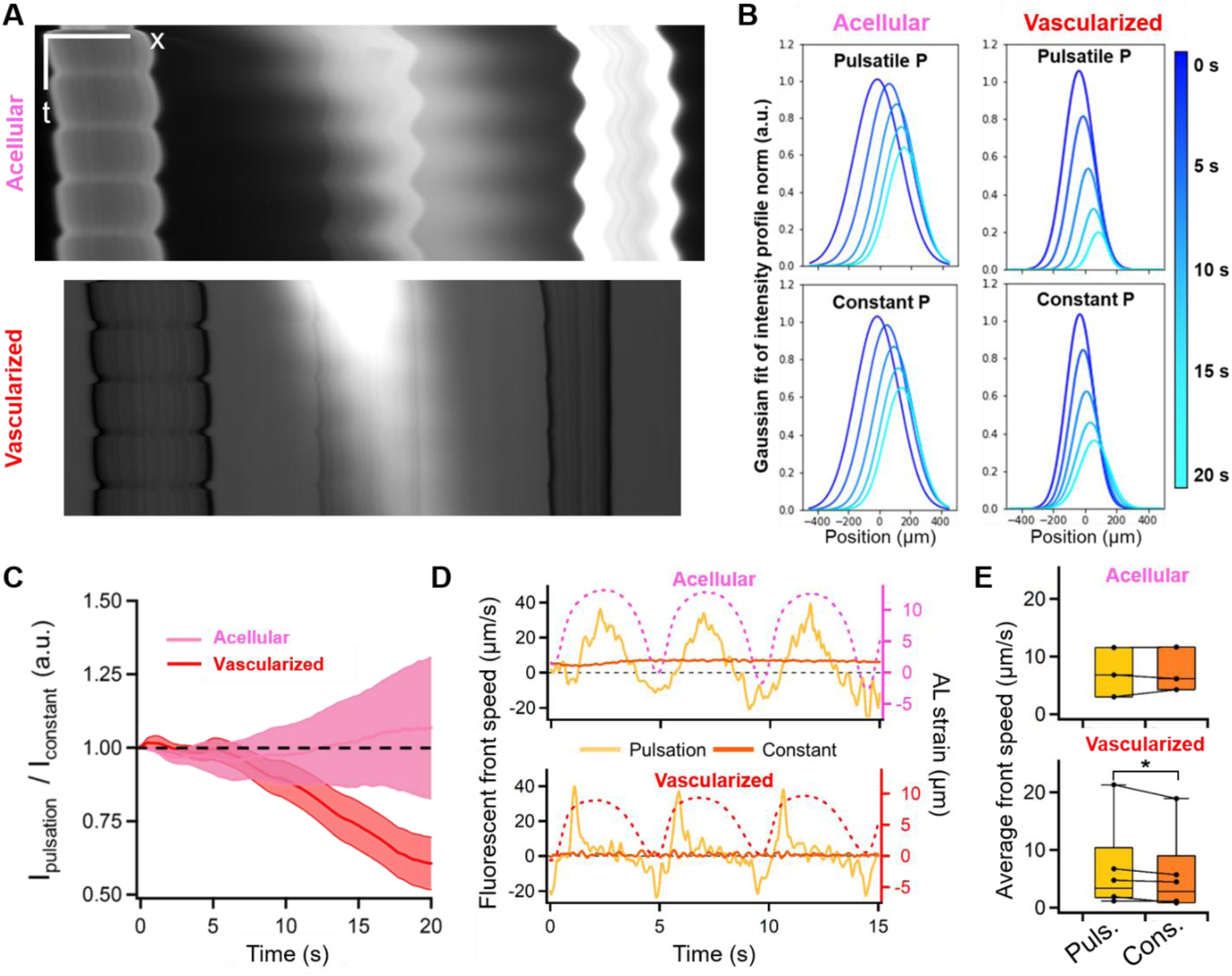
Solute clearance is enhanced by periodic actuation. **(A)** Kymographs of pulsatile movies from acellular and vascularized PIFs. Acellular channels are coated with fluorescent nanoparticles for visualization. Scale bars: 200 µm (horizontal), 5 s (vertical). **(B)** Gaussian fits of the fluorescent signal over time (color scale). Zero coordinate was defined as the CL middle point at t = 0 s. **(C)** Average fluorescence intensity ratio in between lateral lumens (−400 µm ≤ x ≤ 400 µm approx.) under pulsatile vs. constant pressure as a function of time for acellular (n = 3, pink) and vascularized (n = 5, red) PIFs. **(D)** Fluorescent front displacement speed under constant (yellow) or pulsatile (orange) pressure, with AL diameter (dashed lines). Top: acellular PIF; bottom: vascularized PIF. **(E)** Cycle-averaged front speeds under constant and pulsatile pressure. Top: acellular PIF (p > 0.05); bottom: vascularized PIF (p < 0.05).

**Figure 5.**
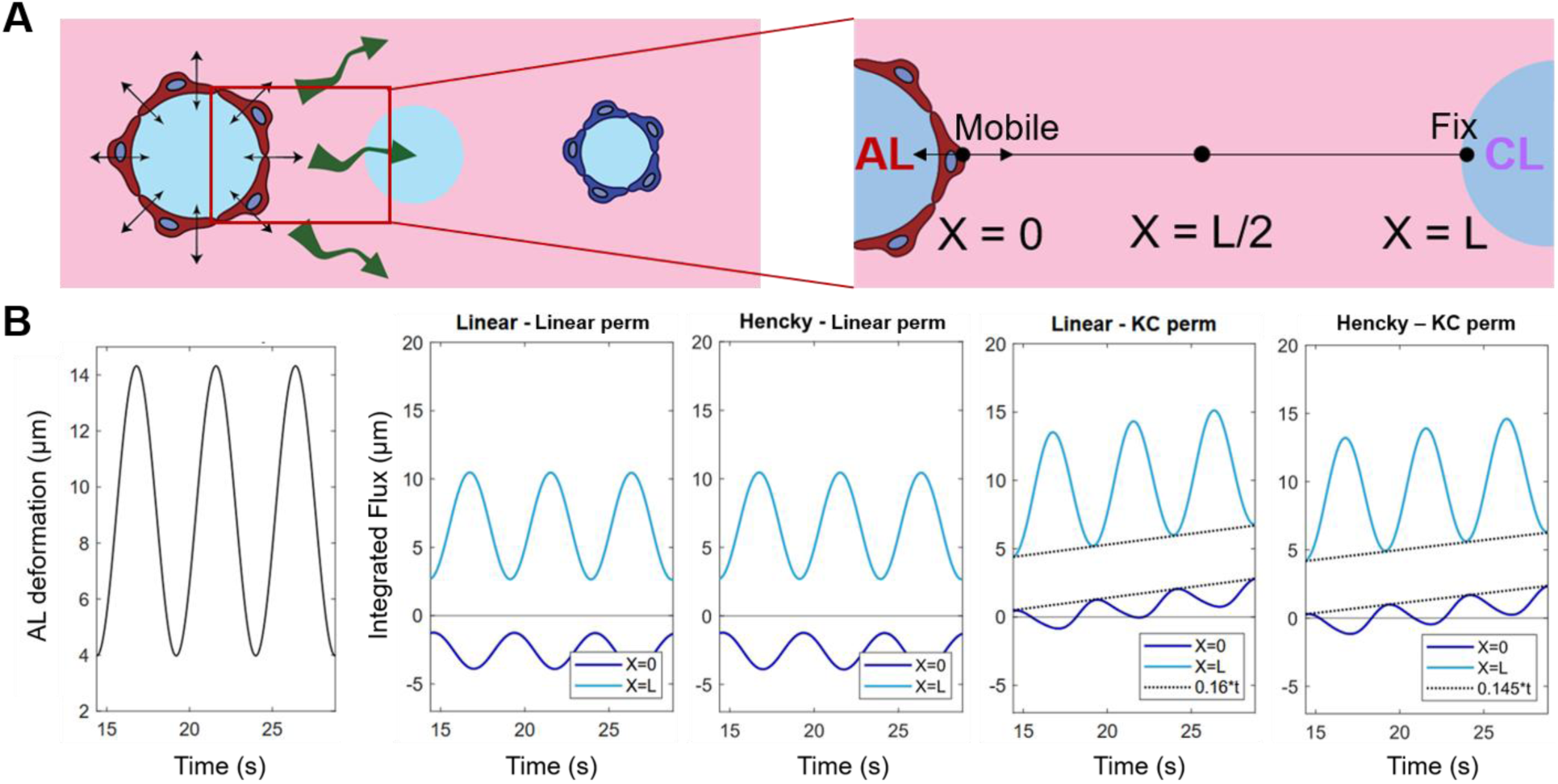
1D poroelastic model of transport. **(A)** Schematic of the 1D AL–CL configuration with boundary conditions (mobile AL, fixed CL). **(B)** Simulations of vascularized 1D model under sinusoidal actuation (black curves) and resulting integrated flux over time at each boundary (light and dark blue curves) for each linear-nonlinear combinations of elasticity and permeability. From left to right: Linear elasticity and linear permeability, Hencky elasticity and linear permeability, linear elasticity and Kozeny-Carman (KC) permeability and Hencky elasticity and Kozeny-Carman permeability. Introducing KC in the 1D model shows a net flux toward the CL, shown as proportional to time at each cycle end (dashed black line).

We characterized the spatial evolution of the dextran concentration profile using Gaussian fitting (Suppl. Fig. 5D), which revealed a time-dependent decrease in concentration amplitude (Fig. 4B). We ruled out photobleaching as a significant factor because we measured a fluorescence reduction of 1% over 20 s in the absence of pressure actuation (Suppl. Fig. 5E). Further, dye observation in the CL lumen (Suppl. Video 5) showed intraluminal transport toward the CL lumen outlets, indicating that fluorescence loss was primarily caused by advective clearance through the CL. We then characterized the contribution of pulsatile pressure actuation to tracer clearance by comparing the temporal variation of the integral fluorescence intensity (numerical integral of the Gaussian fit of the fluorescent signal denoted *I*) under pulsatile *vs*. constant pressure, with the two signals matched in their mean initial value. We computed the ratio *I_pulse_*⁄*I_const_* over time, where a decreasing ratio indicates more effective clearance under pulsatile pressure. Averaging the results obtained with different PIFs, acellular samples showed a constant ratio (*I_pulse_*⁄*I_const_* = 1.05 ± 0.20 after 20 s, Fig. 4C), indicating that the pulsations did not alter the transport of solutes through the collagen scaffold. Conversely, tracer clearance was accelerated under pulsatile pressure in vascularized PIFs (*I_pulse_*⁄*I_const_*= 0.6 ± 0.1 after 20 s), consistent with the lower Gaussian amplitude observed after 20 s under pulsatile compared with constant pressure (0.20 versus 0.36, respectively; Fig. 4B). Thus, intraluminal pulsatile pressure enhanced solute clearance only in vascularized PIFs.

To elucidate the link between enhanced tracer clearance and MV pulsations, we tracked the progression of the fluorescent front in the AL–CL region. We defined the front position as the location where fluorescent intensity was 30% of the maximum, and extracted the front displacement speed by taking the time derivative of the front position and comparing it with AL deformation. In acellular PIFs, the front velocity was highest at the peak of AL expansion (orange curve, Fig. 4D, upper panel), as expected for advection-dominated transport, because the velocity peak corresponds to the maximum pressure difference between AL and CL. Similarly, the front velocity was minimal, and occasionally slightly negative, at the same moment of minimal AL deformation. The cycle-averaged front velocity was similar under pulsatile (6.4 ± 2.7 µm/s) and constant pressure (6.8 ± 1.9 µm/s) conditions (Fig. 4E, top panels). Notably, these velocity values are consistent with Darcy’s law predictions of ∼5 µm/s, taking *κ_gel_* = 10^−1^^4^ m^2^ for the gel permeability, *μ* = 1 mPa·s for the interstitial fluid viscosity, and a pressure gradient of 400 Pa/mm. Collectively, these results demonstrate that solute transport for this molecular weight in acellular PIFs is governed by advection, and that sinusoidal actuation produces transport outcomes equivalent to those obtained under its time-averaged constant-pressure counterpart.

Front progression in vascularized PIFs differed markedly: AL dilation produced a velocity burst, followed by slow front progression for the remainder of the pulsation (Fig. 4E, lower panel), resulting in temporal evolution that was uncorrelated with the sinusoidal pressure input but followed the dynamics of matrix solid deformation. During MV retraction, the front speed became negative, but the absolute velocity burst was lower than during the strain accumulation phase. The cycle-averaged front velocity of 4.3 ± 3.6 µm/s with pulsatile pressure was 16% faster than under constant actuation (3.7 ± 2.7 µm/s; Fig. 5E, bottom panel). This difference in front velocity is consistent with the enhanced clearance observed in Fig. 4C, as the accelerated solute velocity accounts for the clearance in CL. Collectively, solute transport in acellular systems primarily relies on pressure-driven interstitial flow, expectedly resulting in equivalent transport under pulsatile and constant pressure, whereas asymmetric pulsatile strain dynamics in vascularized PIFs are associated with directional solute transport and enhanced clearance.

### 1D poroelastic simulations reveal the role of the barrier’s nonlinear permeability in the enhanced clearance mechanism

Using finite element simulations, we finally investigated the key physical principles accounting for enhanced clearance, such as (i) strain accumulation, (ii) the amplitude and shape of the applied displacement, and/or (iii) the poroelastic constitutive laws describe the MV. We developed a one-dimensional large-deformation poroelasticity model of the AL–CL space, where the vasculature and the collagen are two layers with distinct elastic moduli, permeabilities, porosities, and thicknesses (see Methods, Fig. 5A). For boundary conditions, we clamped the gel at the CL (X=L) and imposed displacements at the AL (X=0) without imposing a hydrodynamic pressure difference, thereby decoupling flow generated by AL wall pulsations from flow induced by a pressure gradient. For the sake of simplicity, we started with sinusoidal displacements to identify the physical mechanisms sufficient to induce bulk flow in the model. As simulation readout, we computed the time-integrated fluid flux across the AL (X=0) and CL (X=L) boundaries (dark and light blue curves, respectively in Fig. 5B and Suppl. Figs. 6 and 7).

Our tests first aimed to check the occurrence of a bulk flow by implementing simulations with nonlinear (Hencky) elastic constitutive law for the solid matrix with nonlinear permeability based on the Kozeny-Carman law^43^. We compared the sinusoidal deformation centered around the equilibrium position to an offset sinusoid to reproduce the experimental strain accumulation (4% maximum, Fig. 3F), and a flattened sinusoidal with the same average compression. Bulk flow towards the CL was detected in every case based on the gradual increase of the integrated flux from period to period at both boundaries. We measured integrated flux rates of 0.13 µm/s for the uncompressed sinusoid; 0.14 µm/s for the compressed sinusoid; and 0.10 µm/s for the flattened, compressed signal (Suppl. Fig. 6), showing consistent conversion of the sinusoidal displacement into a directional fluid flow.

We then deepened our understanding of bulk flow generation by analyzing local porosity variation as a measure of local volumetric strain and local Darcy fluid flux in the compressed sinusoid scenario (Suppl. Fig. 7A, B). In acellular settings, the AL and CL boundaries exhibited equal strains and opposite local fluid fluxes, resulting in symmetrical integrated fluxes at each boundary and therefore no bulk flow. In contrast, the vascularized simulation showed an instantaneous spatial asymmetry with more strain accumulated in the AL than in the CL, as anticipated by our experiments. These simulations also showed slightly unequal influx and efflux locally across the two boundaries, which repeated over multiple cycles, generated a steady fluid flow from the AL toward the CL. Importantly, this linear increase occurred at the same rate at both boundaries, thus maintaining mass conservation in the simulation. In summary, bulk flow arises from the dynamic asymmetries imposed by the vascular barrier over multiple pulsation periods in the nonlinear elasticity, nonlinear permeability 1D model.

Finally, we focused our analysis on how the constitutive laws of vascular barrier poroelasticity impact bulk flow generation. We compared simulation results from linear or nonlinear elasticity, combined with either linear or nonlinear permeability. Linear permeability never produced bulk flow, regardless of the elasticity law implemented, while nonlinear permeability always did (Fig. 5B). These simulations hence identify nonlinear permeability alone, as the constitutive feature required for bulk flow generation in the model. The nonlinear permeability, which captures the coupling between paracellular pore geometry and vascular deformation, is consistent with the observation of strain-dependent microvascular permeability^40^.

Together, these simulations establish a mechanistic route by which MV pulsations can generate directional interstitial flow due to local spatial asymmetries, as experimentally observed. Strain-dependent permeability of the endothelial barrier prevents influx and efflux from cancelling over each cycle, thereby rectifying pulsatile deformation into bulk flow. Within the bilayer 1D poroelastic model considered here, strain-dependent permeability is therefore necessary for the generation of sustained cycle-averaged fluid transport.

## Discussion

In this study, we investigated how MV pulsations modulate solute transport dynamics using an *in vitro* system, the PIF device, which consists of two parallel lumens (both of them either acellular or vascularized) and a tracer injection channel embedded in a collagen matrix. We first characterized the deformation of individual acellular and vascularized lumens, demonstrating that the endothelial barrier localizes hydrogel strain to the immediate vicinity of the MV. We then studied the effects of periodic pressure actuation, observing hyperelastic deformation of the collagen matrix that was more pronounced in vascularized samples, together with greater asymmetry between MV expansion and retraction. We next examined how these pulsatile dynamics affect solute transport. In acellular samples, solute clearance was comparable under constant and periodic pressure actuation, as fluid moved symmetrically back and forth between the AL and PL, without additional net bulk flow beyond that driven by the imposed cycle-averaged pressure gradient. In contrast, vascularization broke this symmetry. MV pulsation produced an increase in fluorescent dextran front velocity under periodic versus constant pressure actuation that accounted for the enhanced solute clearance observed experimentally. We further investigated this response using numerical simulations of flow in a 1D bilayer poroelastic model. These simulations qualitatively reproduced the directional pulsatile fluid motion when strain-dependent permeability was incorporated, where unequal influx and efflux across MV barrier compounded over cycles to produce bulk flow. Based on these findings, we propose a qualitative mechanism of enhanced clearance driven by asymmetric push–refill fluid dynamics (Fig. 6), in which fluid is preferentially displaced toward the CL during MV expansion and replenished from the AL during retraction, thereby generating directional bulk flow over successive pressure cycles.

**Figure 6.**
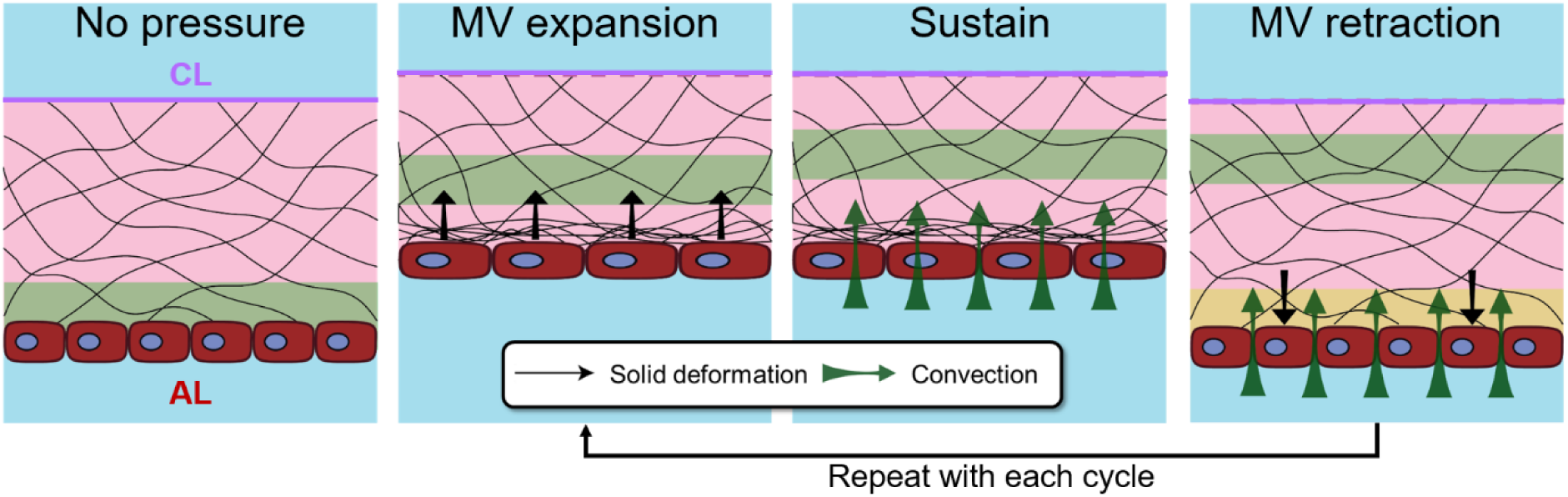
Proposed mechanism of pulsatile deformation-driven transport. Consider a fluid element (green strip) near the AL. Upon pressure actuation, MV expansion is associated with fluid displacement. MV then remains dilated with slow advection toward the CL. Finally, as the MV retracts, fluid is predominantly injected through the MV (yellow strip) rather than the CL to drive a push–refill directional process.

Mechanobiological responses of endothelial barriers to deformation and/or shear stimuli have been widely reported^7,8,46–48^. However, these studies have primarily addressed long-term effects over timescales of minutes to hours, typically assuming that permeability remains constant over shorter timescales, such as seconds. In contrast, our study reveals that vascular permeability is dynamically modulated by strain even in response to small, rapid pressure variations, owing to the deformability of the endothelial barrier. To the best of our knowledge, constitutive laws describing strain-dependent permeability in thin, axisymmetric barrier tissues have yet to be established. This represents both a theoretical and experimental challenge, and highlights the need for new approaches to better characterize and model endothelial barriers in *in vitro* platforms that recapitulate organ-specific mechanical and transport properties. Indeed, liquid permeation through paracellular pores is governed by cell-type-specific expression of junctional proteins ^49,50^ and their interplay with cytoskeletal responses to mechanical stimulation ^48^, thereby highlighting the intrinsically mechanobiological nature of endothelial barrier regulation.

Finally, our findings identify endothelial poromechanics as a potential mechanism for converting vascular pulsations into directional bulk flow with potential implications for tissue fluid and solute clearance. This mechanism may be affected by pathological alterations that increase the stiffness of the extracellular matrix and endothelial tissue, as may occur during aging, potentially impairing pulsation-induced bulk flow and contributing to the loss of tissue homeostasis^51,52^. In the brain, where the glymphatic model of IF circulation has attracted interest, our platform could be further developed to incorporate perivascular cell types, particularly astrocytes, and ultimately enable the self-assembly of a more biomimetic blood–brain barrier model ^53^. More broadly, these findings underscore the need for organ on chip platforms that integrate multiple interacting cell types with instrumentation capable of monitoring dynamic biological functions in biomimetic environments.

## METHODS

### Silicone chip fabrication

Pieces of 3 mm thick clear silicone sheets (As One, Osaka, Japan) were cut using laser cutter (Etcher Laser Pro 30 W, SmartDIYs, Yamanashi, Japan). The device pattern was engraved on the sheets at 100% laser power and a speed of 3500 mm/min in a single pass (Suppl. Fig. 1). The engraved sheets were then cleaned under abundant water with a sponge and soap to remove ash and debris, and immediately dried. The engraved profiles were characterized using a Keyence VHX-7000 microscope (Keyence, Osaka, Japan; Suppl. Fig. 1). Holes for the hydrogel chamber (5 mm diameter) and reservoirs (3 mm diameter) were then manually punched into the engraved silicone pieces using biopsy punchers (Kai Industries, Seki, Japan). The engraved silicone piece was subsequently assembled with a 1 mm thick silicone sheet by exposure to O2 plasma for 1 min in a plasma cleaner (Harrick Plasma, Ithaca, NY, USA). The central cavity was again punched out and the resulting silicone chip was finally bonded to a 32 × 32 mm^2^ glass slide using the air plasma process (Suppl. Fig. 1).

### Cell culture: HUVEC

Chemicals were purchased from Sigma-Aldrich, unless mentioned. Human umbilical vein endothelial cells (Lonza, Basel, Switzerland) were thawed and cultured in 60 mm diameter culture dishes with EGM-2 endothelial cell culture medium (Lonza). HUVECs were expanded and passaged when they reached approximately 90% confluence (at 3-5 days) using 0.25% Trypsin - 0.5 mM EDTA. Culture medium was exchanged every other day and HUVEC culture was kept until passage number 7. HUVECs in suspension were directly injected into the two lateral channels of new PIF chips until complete covering with cells (Suppl. Fig 1) and allowed to mature for 3 days with medium change at the beginning of day 2.

### Double MV fabrication

BSA-soaked acupuncture needles were inserted into the silicone chips. Then, 60 µL of liquid type IA collagen (Nitta Gelatin, Osaka, Japan) was poured into the hydrogel chamber and allowed to reticulate for 10 min at 37°C. After reticulation, the chips, with the needles still inserted, were kept at 37°C, submerged in sterile PBS, for at least one day before use.

For MV fabrication, HUVECs were first harvested from a culture dish and resuspended at a concentration of 4×10^6^ cells/mL in culture medium supplemented with 3% dextran (500 kDa). After the removal of the two lateral needles, the cell suspension was consecutively injected into both MV lumens using a 30G needle to produce a dense cell monolayer (Suppl. Fig. 1). The devices were then incubated for 10 minutes, after which the middle needle was removed, and 0.5 mL of fresh culture medium was added on top of the PIF device. The culture medium was renewed every other day.

To assess barrier quality, we monitored the spatial spreading of fluorescent dextran molecules of 70 kDa for each MV prior to their use in PIF settings (Suppl. Fig 5)^40^. The MV diffusive permeability ranged 2 to 94×10^-7^ cm/s. Whenever required, samples were fixed for 30 min using a para-formaldehyde solution at a concentration of 2% diluted in PBS.

### 3D printed pressure control interface and transport experiment

Pressure interface pieces were first designed using Fusion360 software. The digital models were then 3D printed with an SLA Elegoo Mars 3 Pro 3D printer using clear resin Expert material (W-64C05, NSS Co. Ltd.). The slicing parameters for the pieces included a 50 µm layer height and 2.5 seconds of exposure per layer. To ensure biocompatibility, the 3D-printed pieces underwent secondary curing overnight under a 36 W UV light without thermal treatment, which resulted in total cell viability as confirmed using the Cell Counting Kit-8 (Dojindo Laboratories, Kumamoto, Japan; Suppl. Fig. 2).

This pressure interface was connected to three independent pressure controllers (MFCS, Fluigent, Paris; Suppl. Fig. 2) and mounted on the PIF chip using screws and O-rings to ensure waterproofing. Once sealed, the pressure in all three channels was set and maintained at 100 Pa until the experiments began.

### Microscopies

MV barrier diffusion assays were conducted using confocal laser scanning microscope LMS-700 (Carl-Zeiss, Oberkochen, Germany) at 20× magnification. PIF experiments were recorded with an AXIO observer Z1 microscope (Carl Zeiss, Oberkochen, Germany) paired with a Zyla 5.5 sCMOS high-speed camera (Andor, Belfast, United Kingdom) focusing on the equatorial plane of the MVs at 5× of magnification. MV deformation areas were analyzed to determine the MV diameter on each frame, as described in ref.^48^. The motion of fluorescent tracers was analyzed by a custom-made Python code that averaged and normalized the intensity profile along the recorded area and fitted it with a Gaussian function. Coordinate origin position for the timelapse analysis was set to the center of the frame (not the intensity center) and all fittings, distance and transport tracking was performed according to it. From each Gaussian fit, total dextran signal integral and dextran front position at 30% of the maximum were extracted for intensity analysis and front tracking, respectively.

### COMSOL simulations

We performed finite element simulations of single-lumen poroelastic deformation using COMSOL Multiphysics (COMSOL, Stockholm, Sweden), coupling the Darcy’s Law and Solid Mechanics modules through the Poroelasticity multiphysics interface. A 2D geometry representing a cross-section of the collagen scaffold perpendicular to the lumen axis was simulated. The MV was modeled with an inner diameter of 200 µm and a wall thickness of 3 µm, while the collagen scaffold had a width of 2.5 mm and a height of 1.1 mm. Impermeable and immobile boundary conditions were applied to the silicone and glass walls, whereas the lumen and upper gel surface were considered permeable and deformable. For acellular lumens, two sets of simulations were performed: one assuming linear elastic properties for the collagen gel, with a density of 1000 kg/m³, Young’s modulus of 1 kPa, Poisson’s ratio of 0.3, porosity of 0.99, and permeability of 4 × 10⁻¹⁴ m²; and another incorporating hyperelastic behavior described by the Gent model. The fluid viscosity was always set to 10⁻³ Pa·s in both cases. For MVs, the collagen gel was assumed to exhibit linear elastic behavior, while the endothelial cell layer was modeled as a linear elastic material with a Young’s modulus of 0.7 kPa, a Poisson’s ratio of 0.3, and a parametrizable permeability. The minimum mesh element size was set to 0.3 µm within the endothelial barrier and 5 µm in the bulk collagen gel. The total simulation time was approximately 1 min.

### 1D large-deformation poroelasticity model

The choice behind developing a 1D model is motivated by the requirement of a rigorous large-deformation poroelastic formulation for the specific analysis on the asymmetric poromechanics in a bilayer medium. The geometry represents a single “line” of gel between AL and CL.

To capture the MV – collagen matrix dynamics, we consider a bilayer poroelastic medium of total length *L* = 250*μm*, divided into two layers of lengths *L*_1_ and *L*_2_ respectively, with *L*_1_⁄*L* = 0.05 and *L*_2_⁄*L* = 0.95. Despite corresponding to a thicker barrier than in the real MV-collagen system, this value of *L*_1_⁄*L* ensures computational stability/convergence, and is representative of a generic thin first layer. The values of resting porosity *Φ_f_*_,0_, resting permeability *k*_0_ and p-wave modulus ℳ are 0.2 10^−16^*m*^2^ and 0.7 *kPa* for the MV and 0.99 10^−14^*m*^2^ and 1 *kPa* for the collagen. Note that the reference acellular case is modelled as a homogeneous material with the same properties as collagen. To avoid discontinuities at the interface, each initial property *f*(*X*, 0) — standing for ℳ, *k*_0_ or *Φ*_0_ — is smoothly graded between its two values *f*_1_ and *f*_2_ through a hyperbolic tangent profile,

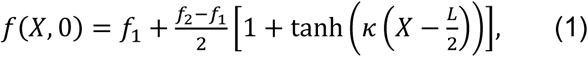

where the steepness coefficient is fixed to *κ* = 40.

The poromechanics model follows a classical formulation for large-deformation poroelasticity^54^. Given the presence of heterogeneities, the model is formulated in Lagrangian coordinates, accordingly to the equations proposed in Fiori and Lorthois^55^ and briefly summarized below. Given the Lagrangian solid displacement field **U_s_**(**X**, *t*) = **x**(**X**, *t*) − **X**, with **x**(**X**, *t*) the current position of the material point whose reference position is **X**, we take the initial, reference configuration to be the relaxed one, so that **U_s_**(**X**, 0) = 0. In 1D and for a uniform initial porosity *Φ_f_*_,0_, the Jacobian determinant *J* = 1 + *∂U_s_*/ *∂X* links the local change in volume to the change in porosity as *J*(*X*, *t*) = 1 − *Φ_f_*_,0_ + *Φ_f_* ⇒ *∂U_s_*/ *∂X* = *Φ_f_* − *Φ_f_*_,0_, where the nominal porosity *Φ_f_* = *Jφ_f_* measures the current fluid volume per unit reference total volume and is expressed as a function of the true porosity *φ_f_*.

Through Darcy’s law, we define 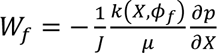 as the Lagrangian nominal flux of fluid through the solid skeleton. With *k*(*X*, *φ_f_*) the deformation-dependent permeability, *μ* the fluid viscosity, and *p* the pore pressure. We take the permeability to follow a normalized Kozeny–Carman relation and the elasticity law to be Hencky elasticity. Combining mass conservation, Darcy’s law and mechanical equilibrium, we derive the final nonlinear flow equations:

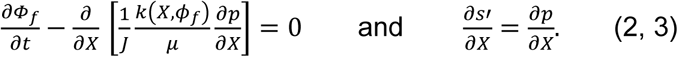

where *s*′ is the effective solid stress. We summarize here the initial and boundary conditions used in this model configuration.

For the initial conditions, we take the left and right boundaries of the solid to be at *X* = 0 and *X* = *L*, with initial conditions *Φ_f_*(*X*, 0) = *φ_f_*_,0_ and *U_s_*(*X*, 0) = 0, *W_f_*(*X*, 0) = 0. At the left boundary we apply the periodic displacement *a*(*t*) derived from experimental data, by imposing *v_s_*(0, *t*) = ȧ(*t*) and *p*(0, *t*) = 0. The right boundary is permeable but fixed in place, so that *v_s_*(*L*, *t*) = 0 (implying *W_f_*(*L*, *t*) = *q*(*t*)) and *p*(0, *t*) = *p*(*L*, *t*), or *s*^′^(0, *t*) = *s*^′^(*L*, *t*)).

For the nondimensionalization, we consider as our reference the acellular case (see properties listed above). Hence, the reference poroelastic timescale used in all simulations is *T*_pe_=(*μL*^2^)/(*k*_0_ℳ), where *μ* is the viscosity of the fluid (water), *L* the total length of the material (unchanged between acellular homogeneous and MV bilayer case) and *k*_0_ and ℳ are the initial permeability and p-wave modulus of the collagen gel. The equations are then scaled by 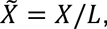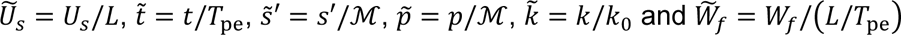.

We solve the model numerically in MATLAB, using compact finite differences in space with implicit Runge–Kutta integration in time, via the built-in solver ODE15s (analogously to Fiori et al., 2023)^56^.The convergence study for the model is proposed in Suppl. Fig. 7B,C.

## Supporting information

Pulsating acellular PIF fluorescent video

Pulsating vascularized PIF fluorescent video

Constant pressure in acellular PIF fluorescent video

Constant pressure in vascularized PIF fluorescent video

Central lumen fluorescent evacuation in pulsating vascularized PIF video

## Acknowledgements

This research was partly supported by WINGS-QSTEP (D.A.), LIMMS, and ARIM of the MEXT, Grant Number: JPMXP1223UT-0331 and by a grant from the Leducq Foundation and the Leducq Foundation for Cardiovascular Research (23CVD03). The authors thank Dr. Vincent Salles for his contributions on the use of the Keyence VHX-7000 microscope.

## Author contribution declaration

D. Alcaide Martín carried out the experiments and data analysis under the supervision of Y.T. Matsunaga, J. Cacheux and A. Bancaud. D. Alcaide Martín and J. Cacheux conceptualized the device and fabrication process. A. Bancaud and J. Cacheux developed the numerical simulations supporting the experimental work, while M. Fiori and S. Lorthois conceptualized and carried out the theory-based poroelastic simulations. All authors revised, corrected and contributed to the writing of the manuscript, mainly produced by D. Alcaide Martín.

## Data availability statement

The data that support the findings of this study are openly available in a Figshare repository at https://doi.org/10.6084/m9.figshare.31933326.

