## Supplementary figures and Supplementary video captions for "Pulsations of engineered microvessels enhance solute clearance via heterogeneous strain-dependent poroelasticity"

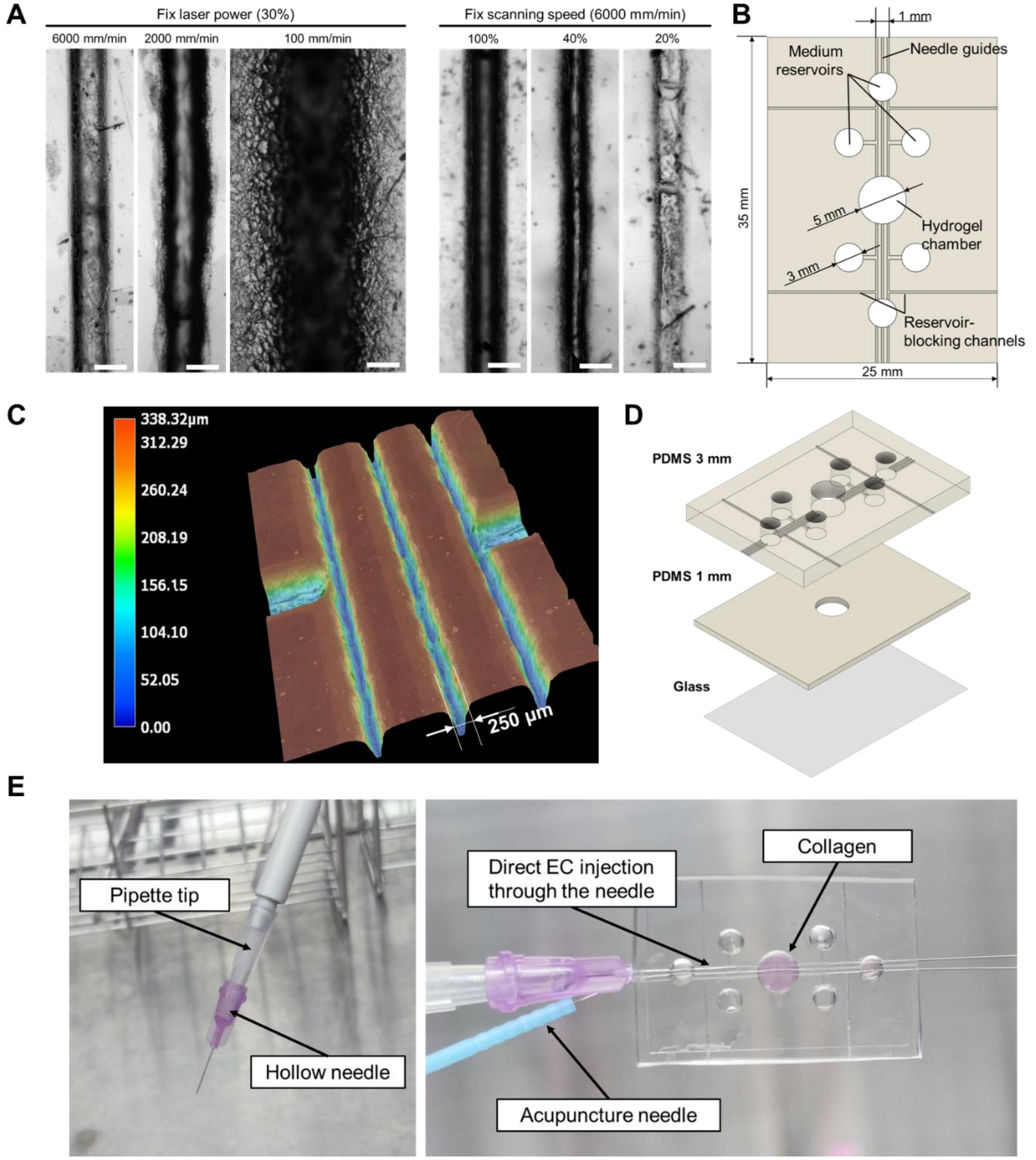


**Supplementary Figure 1 – (A)** Grooves created through different power and speed settings of the Laser engraving machine. **(B)** Detailed dimensions of the PIF chip design to be replicated by laser engraving over PDMS sheets. **(C)** Optical profile of laser engraved central channels and the reservoir connection channels. 3500 mm/min at 100% of power produce U-shaped grooves 250μm wide and 340 μm deep, suitable for sliding 200 μm diameter acupuncture needles through them. **(D)** Layers plasma bonded to seal and finalized PIF device fabrication. **(E)** Pipette tip modification to fir a 30G tube needle for cell injection in the MV channels.

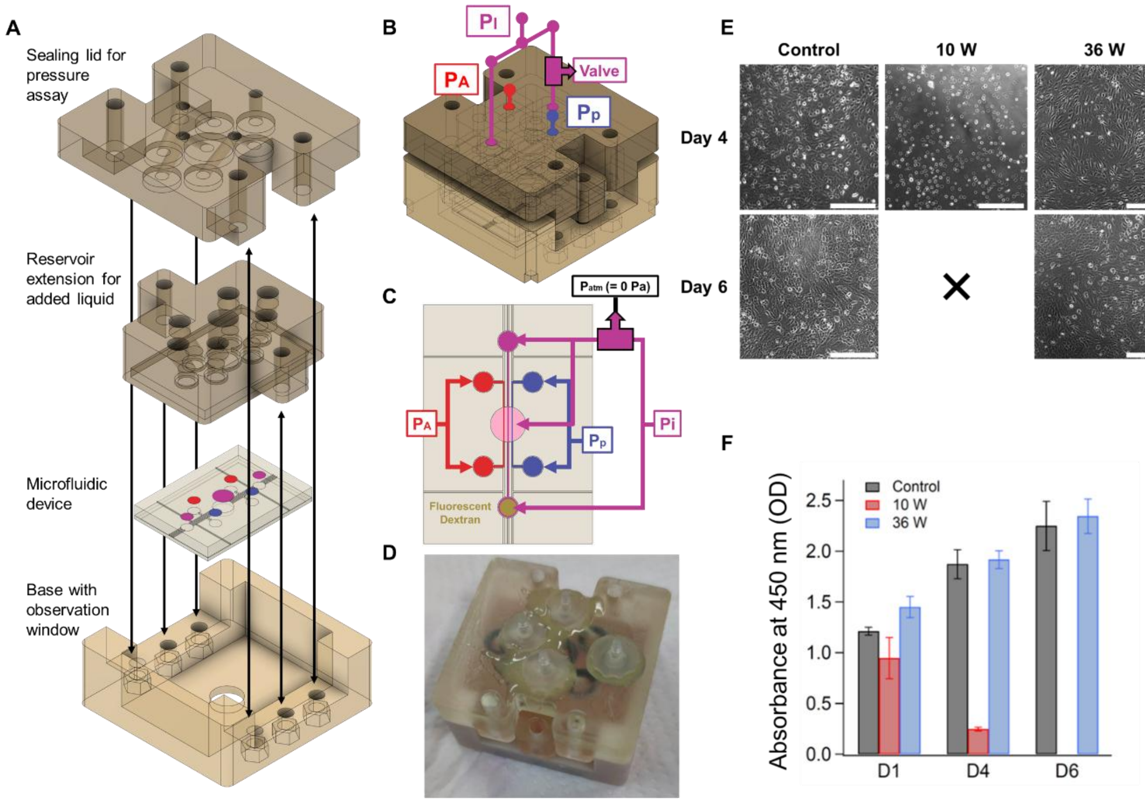


**Supplementary Figure 2 – (A)** 3D models of the pressure interface pieces. Device sealing is achieved by tightly screwing all pieces together. **(B)** 3D model of the complete pressure interface assembly with pressure connections indicated by color: Red for Actuation pressure [$P_A$], purple for injection pressure [$P_I$] (with a manual valve in one side) and blue for Passive pressure [$P_P$]. **(C)** Pressure input into the PIF device schematic. Fluorescent dextran solution is injected in the injection channel reservoir opposite to the valve, so that when opened, flow in the injection channel drags tracers along into the collagen. **(D)** Picture of the closed pressure interface of the PIF device. **(E)** Phase contrast images of HUVEC cell culture in wells where 3D printed pieces post-treated under 10 W or 36 W UV lights overnight were submerged. Pieces treated at 10 W produced generalized cell death by day 4 of culture. **(F)** Cell-counting kit 8 absorbance measurements for wells containing 3D printed pieces. Absorbance increase is related to cell growth. Control and 36 W treated wells show similar cell growth dynamic.

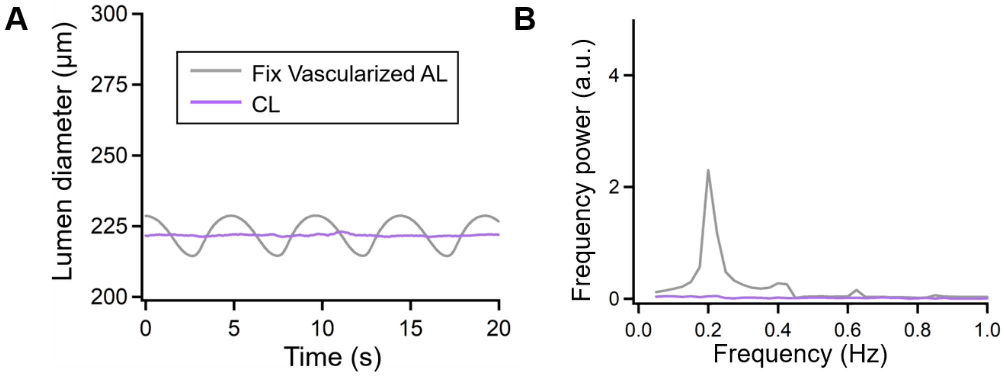


**Supplementary Figure 3 – Fix vasculature lumen response. (A)** AL and CL diameter response under pulsatile pressure after PFA fixing. **(B)** Fourier transform of each diameter signal.

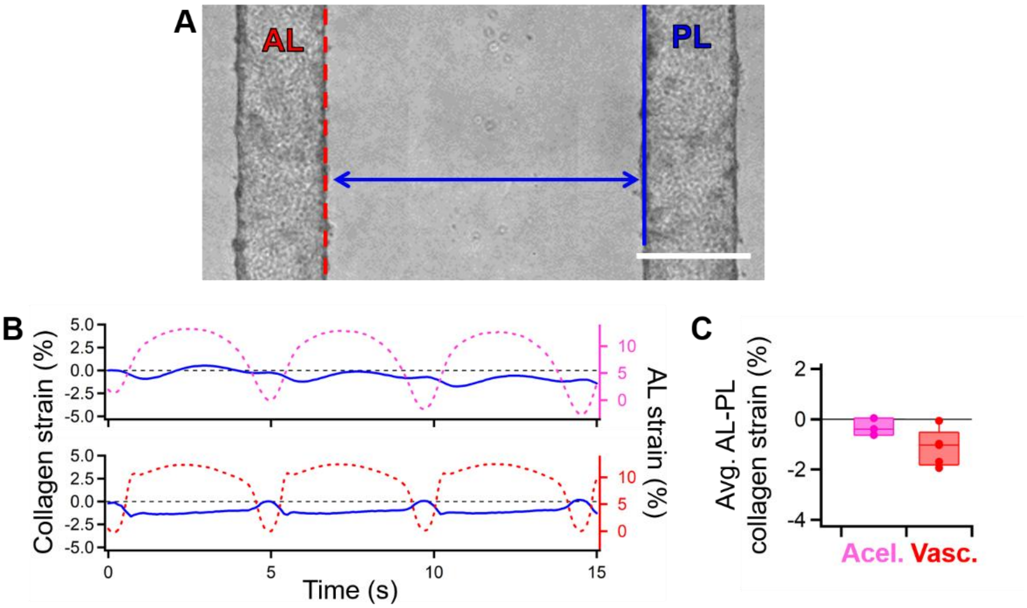


**Supplementary Figure 4 – AL-PL collagen deformation response. (A)** Depiction of AL-PL distance tracking strategy (blue arrow). Scale bar = 300 µm. **(B)** AL-PL collagen gel strain obtained from inter-lumen distance variation (blue curves) under pulsatile actuation: acellular PIF (top) and vascularized PIF (bottom). Dashed curves represent AL strain over time, as deduced from Fig. 3. **(C)** Cycle-averaged AL-PL collagen strain in acellular (n = 3) and vascularized (n=5) PIFs ($p > 0.05$).

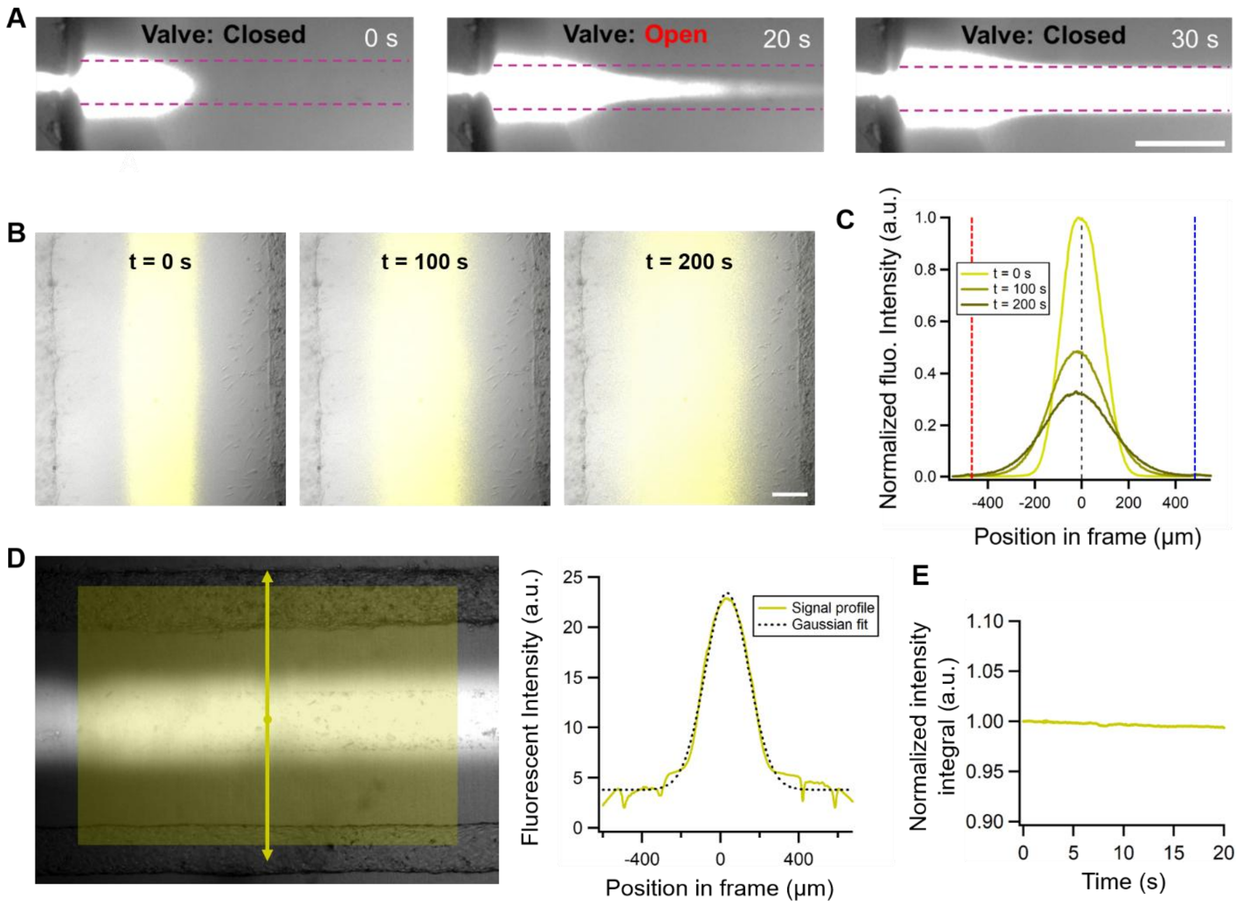


**Supplementary Figure 5 – Fluorescent dextran injection and signal analysis in PIF. (A)** Dextran injection by opening the valve on the right side of the PIF and creating a 100 Pa pressure difference between the CL extremes. **(B)** Confocal fluorescent images of 70 kDa FITC-dextran diffusion from the CL overlayed on PIF bright field image. AL and PL borders can be seen in the frames, which are barely reached by the dextran after 200 s. **(D)** Confocal fluorescent dextran diffusion profiles. AL and PL borders are represented as red and blue dashed lines, respectively. Dextran spreading corresponded to 52 $\mu m^2/s$ diffusion coefficient, calculated through gaussian fitting of the signals. **(E)** Left - Wide view of dextran-filled CL in vascularized PIF. The intensity profile analyzed is obtained in the direction of the yellow arrows and averaged in the shaded area. Right - Intensity profile of image B. The fluorescence intensity is off-set by the background light of the image. Gaussian fit of the data is shown. **(D)** Normalized fluorescent intensity integral over time in the absence of actuation pressure.

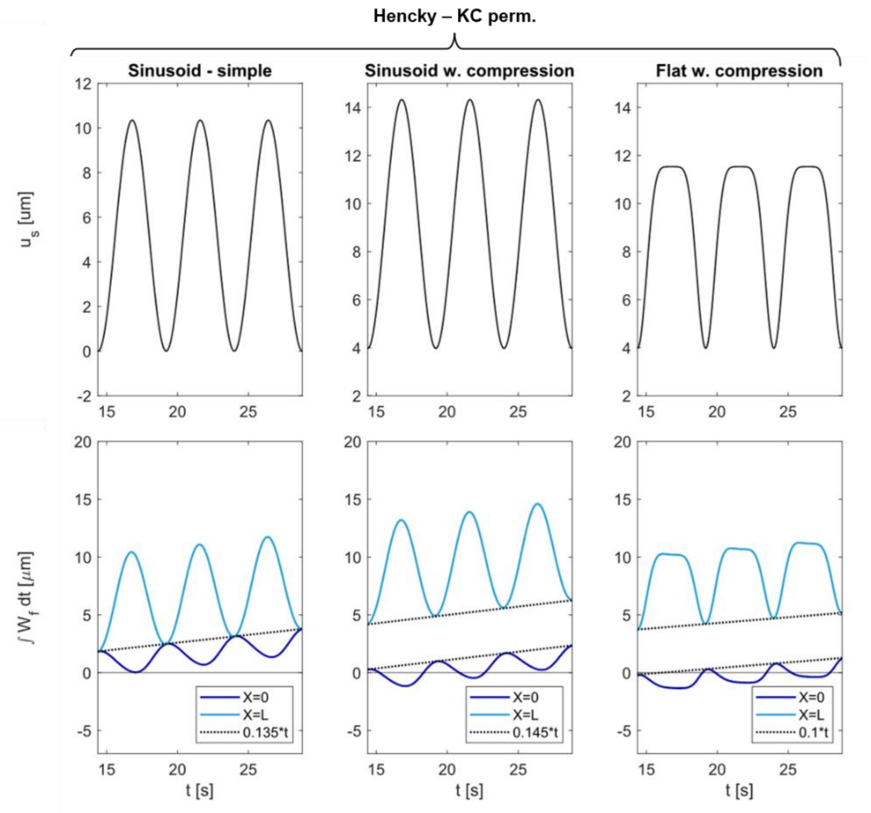


**Supplementary Figure 6 - 1D model simulation with different waveforms**. Simulations of vascularized 1D model under different actuation (black curves), and resulting integrated flux over time at each boundary (light and dark blue curves). Simulations with Henky elasticity and Kozeny-Carman permeability, from left to right: sinusoidal deformation with no strain offset; sinusoidal deformation with strain offset; flattened sinusoidal deformation (maintaining same strain average than normal sinusoidal input) with strain offset. All of them produce bulk flow.

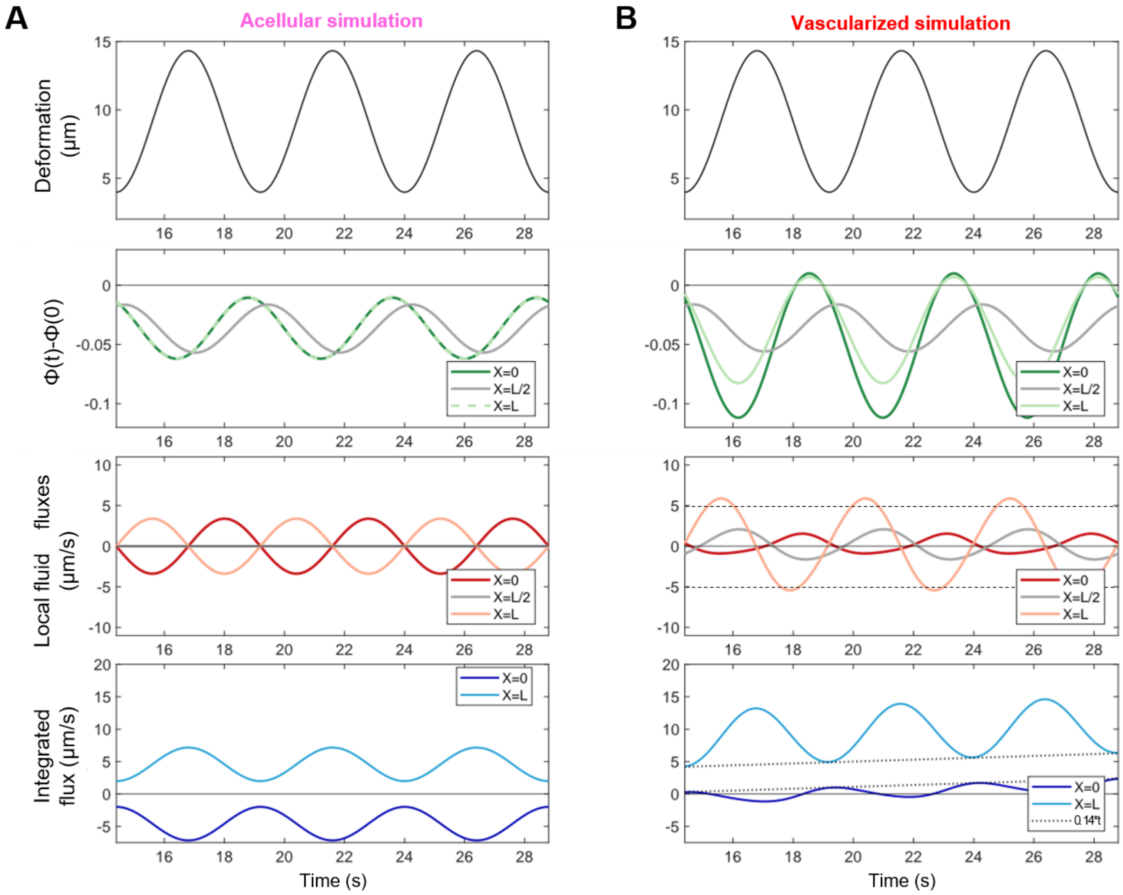


**Supplementary Figure 7 –Acellular vs Vascularized 1D simulations. (A)** Acellular simulation inputs and results. The first row of panels shows deformation graph: compressed sinusoid actuation (black curve). The second and third rows report porosity variations (green) and local fluid fluxes (red) at X=0, L/2, and L with shaded colors, respectively. The fourth row summarizes the integrated flux at each boundary. No bulk flow is reported. **(B)** Vascularized simulation inputs and results, same as (A). More compression is observed in the AL boundary (X=0) and local flows are asymmetric in both boundaries, favoring flow in the CL direction (positive flux). This local imbalance results in bulk flow appearing (dotted black lines in Integrated flux plot).

### Supplementary Videos

**Supplementary Video 1 –** Fluorescent dextran displacement in acellular PIF under pulsatile pressure. The active channel (left) swells the collagen hydrogel, displacing laterally the central and passive channels. Fluorescent dextran is observed to penetrate the collagen region between the central and passive channels, indicating the existence of flow in this collagen region apart from ejecting flow through the central channel (hence the decrease in fluorescent intensity).

**Supplementary Video 2 –** Fluorescent dextran displacement in vascularized PIF under pulsatile pressure. The active vessel (left) expands and retracts sharply without greatly displacing the other two channels. Fluorescent dextran is observed to remain at the central channel collagen border, probably due to blockage of any flow towards the passive vessel by its cellular barrier. This results in most of the dextran to evacuate the ROI through the central lumen.

**Supplementary Video 3 –** Fluorescent dextran displacement in acellular PIF under constant pressure. The active channel (left) swells the collagen hydrogel and displaces the other two channels until the pressure gradient between them is established. Fluorescent dextran is observed to smoothly travel towards the passive channel at a similar rate than in the pulsating acellular case.

**Supplementary Video 4 –** Fluorescent dextran displacement in vascularized PIF under constant pressure. The active vessel (left) expands quickly and fluorescent intensity is observed to decrease gradually, without much lateral displacement, most likely due to the ejection of molecules through the central channel. Compared to the vascularized pulsatile case, dextran ejection rate appears slower.

**Supplementary Video 5 –** Visualization of dextran ejection through the central channel in vascularized PIF device. Fluorescent signal, contrast, and color scheme have been pos-processed for better visualization of the dextran waves that occur in the central channel (slightly darkened in this movie due to some early dextran ejection). From the two bright vertical dextran fronts at each side of the central lumen, the left one, closer to the active MV, suffers wave-like motions in the vertical direction of the video, proving the creation of flow in the central channel perpendicular to the pressure actuation gradient.
